# The Q226L mutation can convert a highly pathogenic H5 2.3.4.4e virus to bind human-type receptors

**DOI:** 10.1101/2025.01.10.632119

**Authors:** María Ríos Carrasco, Ting-Hui Lin, Xueyong Zhu, Alba Gabarroca García, Elif Uslu, Ruonan Liang, Cindy M. Spruit, Mathilde Richard, Geert-Jan Boons, Ian A. Wilson, Robert P. de Vries

## Abstract

H5Nx viruses continue to wreak havoc in avian and mammalian species worldwide. The virus distinguishes itself by the ability to replicate to high titers and transmit efficiently in a wide variety of hosts in diverse climatic environments. Fortunately, transmission to and between humans is scarce. Yet, if such an event were to occur, it could spark a pandemic as humans are immunologically naïve to H5 viruses. A significant determinant of transmission to and between humans is the ability of the influenza A virus hemagglutinin (HA) protein to shift from an avian-type to a human-type receptor specificity. Here, we demonstrate that a 2016 2.3.4.4e virus HA can convert to human-type receptor binding via a single Q226L mutation, in contrast to a cleavage-modified 2016 2.3.4.4b virus HA. Using glycan arrays, x-ray structural analyses, tissue- and direct glycan binding, we show that L133aΔ and 227Q are vital for this phenotype. Thus, whereas the 2.3.4.4e virus HA only needs a single amino acid mutation, the modified 2.3.4.4b HA was not easily converted to human-type receptor specificity.

## Introduction

Influenza A viruses (IAV) natural hosts are wild waterbirds, from which they can cross the species barrier to gallinaceous poultry and, albeit rarely, to various mammalian species (1). IAVs carry two protruding envelope glycan proteins, in which the hemagglutinin (HA) binds sialic acids on the cell surface, and the neuraminidase (NA) cleaves them to enable progeny escape from infected cells. Antigenically, IAVs are clustered by their HA and NA proteins, of which we now have 17 and 11 subtypes in avian species, respectively, as well as two others in bats (2). While 17 HA subtypes can infect birds, and only a few viruses have crossed the species barriers to mammals, for example, H3N8 in horses, H1N2 in pigs, and H1N1, H2N2, and H3N2 in humans. The recent outbreak in dairy cattle in the USA now represents a large mammalian reservoir from which H5N1 viruses can infect humans and possibly adapt (3).

H5Nx viruses, where x stands for different NA possibilities, are one of the two subtypes, together with H7Nx, that can be highly pathogenic in chickens (4). This high pathogenicity is due to a multibasic cleavage between the HA1 and HA2 subunits of HA, leaving the HA susceptible to host furin-like proteases. Since the emergence in 1997 of the goose/Guangdong strain (5), H5 influenza viruses have infected more than 900 humans, with an overall fatality rate of 50% (6, 7). Fortunately, H5Nx viruses are unable to sustainably transmit between humans. A primary barrier is the inability of these viruses to efficiently bind human-type receptors (8). Human-type receptors are defined by a sialic acid (Sia) α2,6-linked to a galactose, which in turn is β1,4 linked to an N-acetylglucosamine (LacNAc). Instead, H5 influenza viruses maintain their binding preference to avian-type, α2,3-linked Sia. The mutational pathways to achieve human-type receptor binding for H5 IAVs have been described in detail and differ for different genetic and antigenic backgrounds (8).

The classical amino acids conferring human-type receptor specificity are G225D and E190D for H1 and other group 1 influenza viruses, while Q226L and G228S are essential for H3N2 and other group 2 viruses (9–11). In previous studies, H5N1, although group 1, appears to need Q226L as well as other mutations to bind human-type receptors (8). Some of the authors in this study recently demonstrated that a A/Texas/324/24 H5 HA protein, a human isolate from the current 2.3.4.4b outbreak did gain human-type receptor binding when Q226L was introduced (12). However, in earlier H5 HAs, additional mutations, such as R/K193S/T/M, Q196R, N224K, and G228S are often required to bind α2,6 linked Sia exclusively, since single mutations often retain avian-type receptor specificity or result in null mutants. In fact, in the majority of HA subtypes, multiple amino acid changes are required for optimal binding to human-type receptors (8, 13–23). Therefore, every genetic or antigenic strain must be evaluated at the molecular level to determine which amino acid changes confer human-type receptor binding as the same mutation can be highly context dependent.

Since late 2020, we have been experiencing an unprecedented global outbreak of highly pathogenic H5Nx influenza A viruses (IAV) (24). These viruses cause incredibly high mortality in avian species with significant transmission to mammals (25–28) and are circulating year-round from the Northern to Southern hemisphere. With such a bevy of highly pathogenic viruses circulating, resulting in distinct antigenic groups, it is vital to establish the molecular determinants for switching from avian- to human-type receptor binding specificity. Currently, circulating 2.3.4.4b viruses retain avian-type receptor specificity (29–31) and recent literature demonstrated that the Q226L is indeed crucial for infection of cells that display human-type receptors (12, 32). However, molecular details on different receptor binding platforms is lacking. We demonstrate that an H5N6 A/black swan/Akita/1/2016 HA that represents a minority variant 2.3.4.4e (33), containing two distinct amino acid changes in the receptor binding site (RBS) (L133aΔ (i.e. no residue L133a) & R227Q), only requires a single Q226L mutation to bind human-type receptors specifically. On the other hand H5N8 A/duck/France/1611008h/16 HA (with a monobasic HA1/HA2 cleavage site), which is in the same clade as the currently dominant 2.3.4.4b strain, was not converted to bind linear human-type receptors.

## Results

### H5Nx 2.3.4.4b and e antigenic clades are not equally well converted to bind human-type receptors

With continued H5Nx outbreaks in avian and mammalian species, we focused on comparing a model H5 derived from H5N1 A/Indonesia/5/05 (Indo05), which we have studied extensively (34), with two antigenically distinct early H5N8 and H5N6 2.3.4.4 viruses representing the dominant b clade (A/duck/France/1611008h/16) (France16) and an e clade (A/black swan/Akita/1/2016) (Akita16) (35). Whereas the Indo05 HA differs extensively in and around the receptor binding domain, the HA of France16 only differs at six amino acids positions (L111M, L122Q, T144A, T199I, V214A, and N240D) in HA1 from the H5N1 that is in circulation now in cattle in the USA (Fig S1). To assess receptor binding specificity, we used a miniaturized glycan array using a set of biantennary N-glycans with 1 to 3 LacNAc repeats terminating with galactose, α2,3 and α2,6 Sia respectively (Fig 1A) (36). We employed this set of N-glycans as these are biologically relevant structures found in the avian and human respiratory tract (37, 38). The model H5 of Indo05 solely bound to the avian-type receptors depicted in white bars, and the Q226L mutation abrogated all binding (Fig 1B) (34). Next, we tested an H5N8 A/duck/France/161108h/2016 (a kind gift from Romain Volmer, Universite Veterinaire de Toulouse, France), that was intended to represent the contemporary circulating and dominant 2.3.4.4b clade family. This wild type HA protein only binds avian-type receptors (Fig 1C); when Q226L is introduced strong binding is still observed in this assay for avian-type receptors, but with some weak binding for human-type receptors. When we added G228S, which commonly co-evolves with Q226L, this H5 HA protein did not show any responsiveness to any glycan on the array (Fig 1C). Finally, we repeated the same experiment with the 2.3.4.4e clade, which contains two distinct amino acid changes in the receptor binding site (L133aΔ & R227Q). A/black swan/Akita/1/16 (Akita16) (a kind gift of Masatoshi Okamatsu, Hokkaido University, Japan) H5 solely bound to avian-type receptors (Fig 1D); surprisingly however, the Q226L mutation entirely conferred binding to human-type receptors. Adding G228S did not change this phenotype and somewhat reduced responsiveness to the array. Due to the surprising ease of converting Akita16 with a single Q226L mutation to bind human-type receptors, we reintroduced the aΔ133L and R227Q into the Akita16 2.3.4.4e receptor binding site (Fig 1E). This protein bound only to avian-type receptors, and now, with the introduction of the Q226L, bound very weakly to any glycan on the array. We finally used an H3 protein derived from A/Netherlands/761/2009 as a human-type specific control (Fig 1F). We conclude that the 2.3.4.4e receptor binding site conformation is vital to quickly adapt to human-type receptors. Importantly the introduction of Q226L in three related yet antigenically distinct H5 HA proteins led to three different phenotypes, ranging from little change in avian-type specificity, to no binding to avian- or human-type receptors, or complete reversion to human-type receptor specificity.

**Figure 1.**
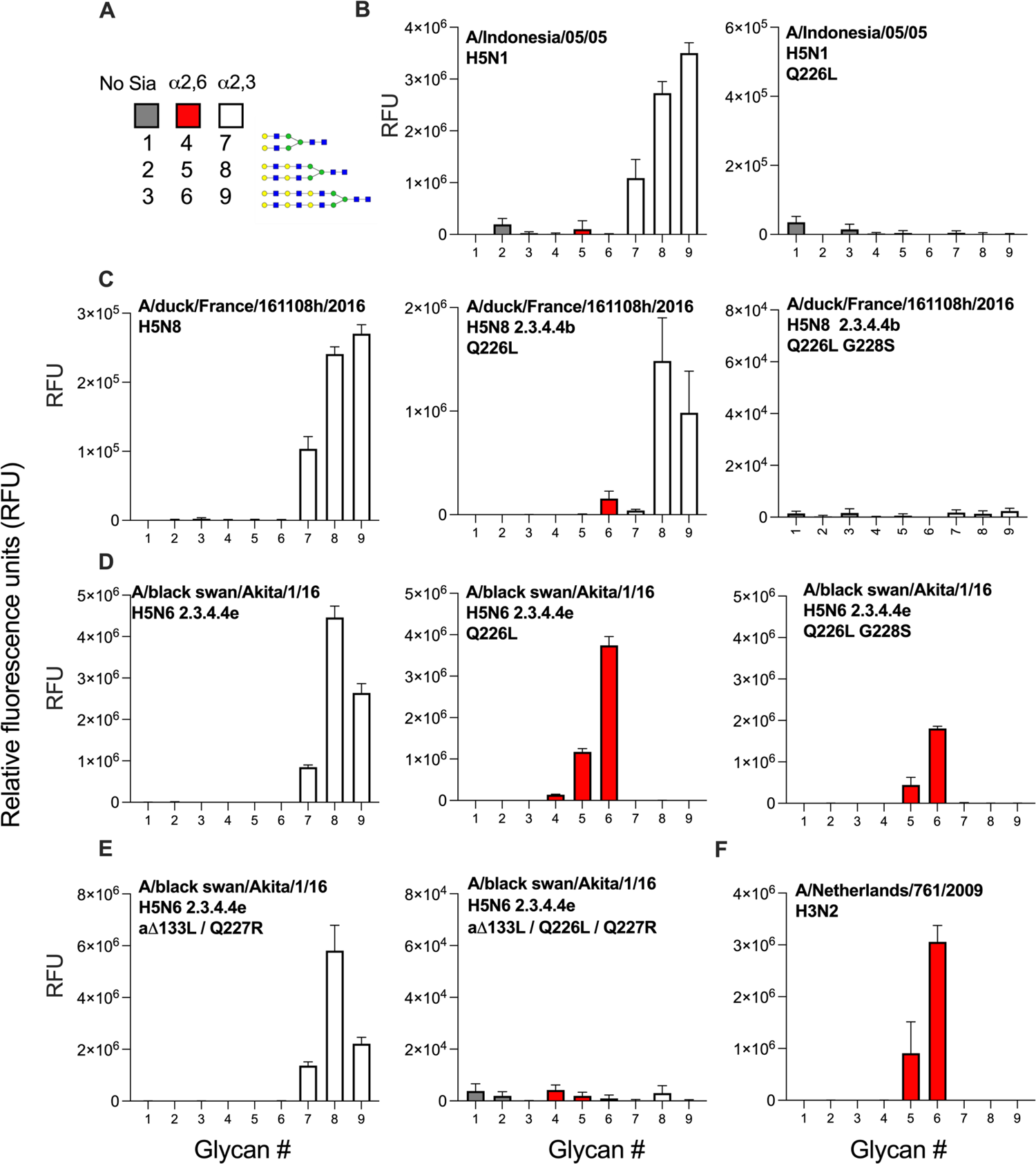
Glycan microarray analyses of H5 hemagglutinins including Q226L and G228S mutants. Synthetic bi-antennary N-glycans printed on the microarray, either without sialic acid (structures 1-3, gray), with α2,6-linked NeuAc (4-6, red), or α2,3-linked NeuAc (7-9, white). Structures 1, 4, and 7, contain one LacNAc repeat, while structures 2, 5, and 8 have two repeats and structures 3, 6, and 9 contain three repeats (A). Indo05 and the Q226L mutant (B). France16 2.3.4.4b as wt, with the single Q226L and G228S added (C). Akita16 as wt, with Q226L and G228S added (D). Akita16 with the 2.3.4.4b specific mutations L133a & R227Q, and Q226L added (E). A/Netherlands/761/09 as the human-type receptor specific control (F). Bars represent the mean ± SD (*n*= 4)

**Figure 2.**
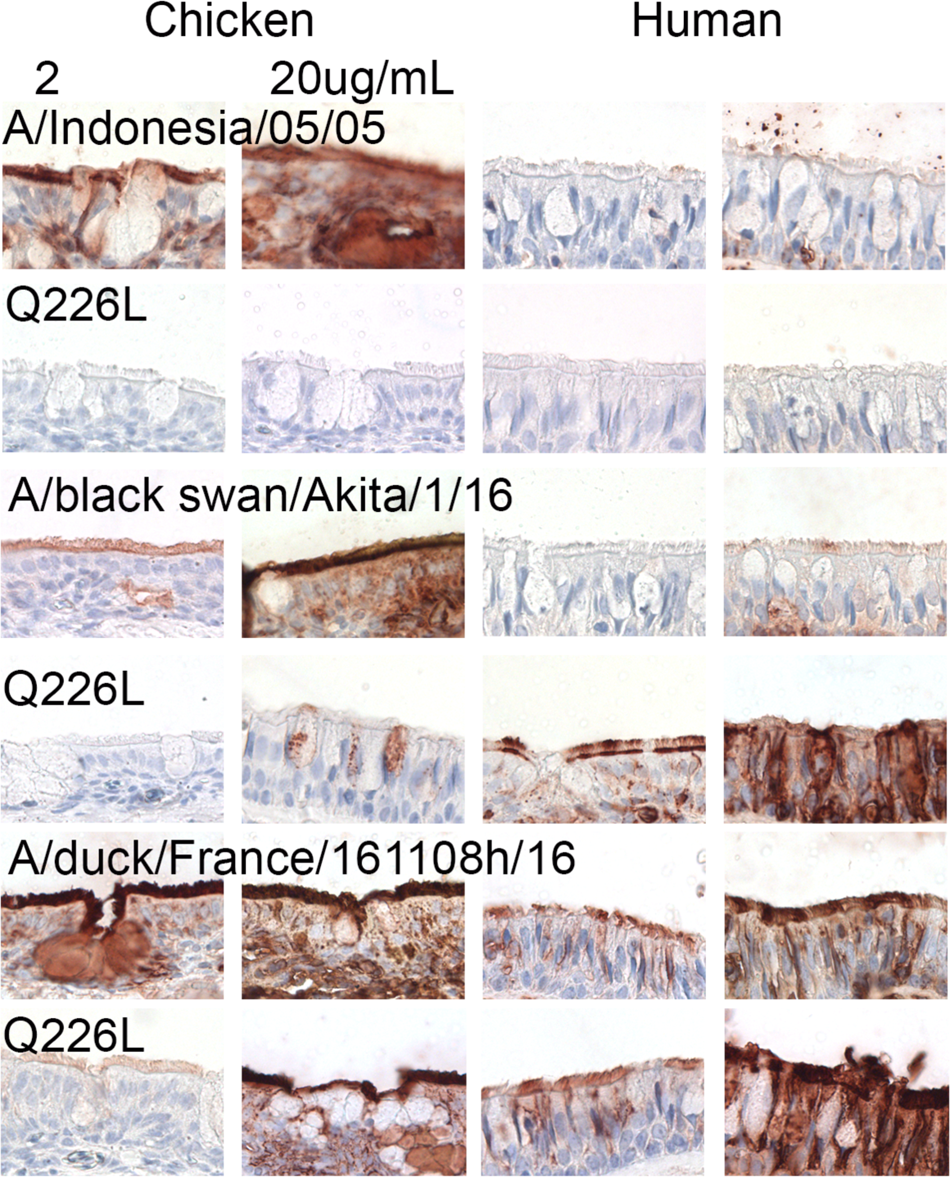
Tissue binding properties of different H5 proteins on chicken and human tracheal slides reveal different phenotypes conferred by Q226L. The binding to chicken and human tracheal tissue was investigated for different influenza A H5 HAs at two different concentrations. From top to bottom Indo05, Akita16 and France16 including their Q226L mutant. AEC staining was used to visualize tissue binding.

**Figure 3.**
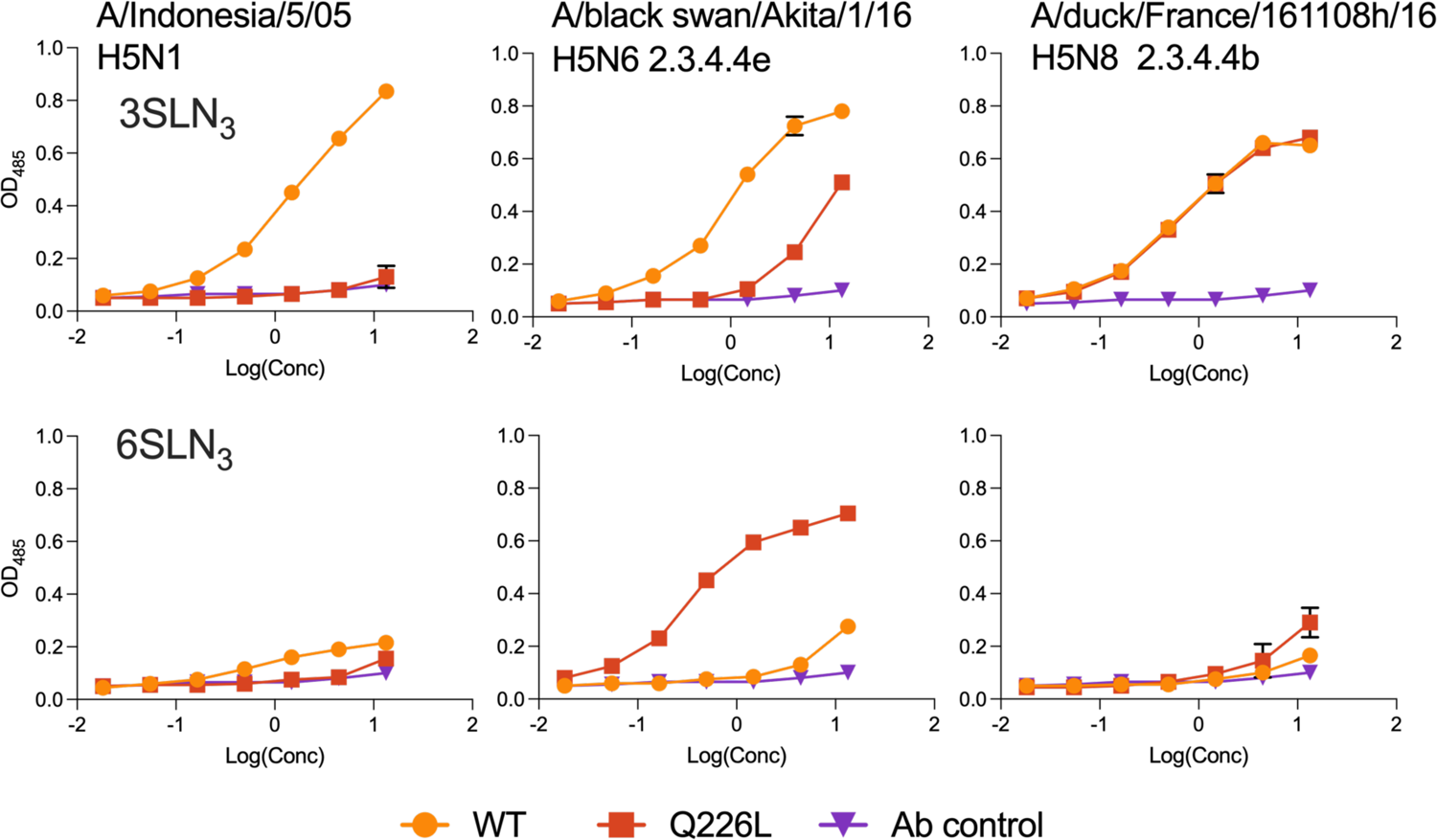
Direct receptor binding assay using biotinylated linear avian- and human-type receptors. 3SLN_3_ (top) and 6SLN_3_ (bottom) coated streptavidin plates interrogated with A/Indonesia/05/05 (left), A/black swan/Akita/1/16 (middle) and A/duck/France/161108h/16 (right) WT proteins as shown with orange circles and the Q226L mutants using red squares.

### The Q226L mutation in A/black swan/Akita/1/16 also confers binding to human upper respiratory tissues

The ability of IAVs to bind either cell in the chicken or human upper respiratory tract is a strong proxy for the ability of a virus to infect either species. Here, we employed the identical proteins as in the glycan array and subjected them to chicken and human tracheal sections at low (2 μg/mL) and high (20 μg/mL) protein concentrations. The H5 protein from Indo05 solely bound to the chicken tracheal sections and, even at a high concentration, failed to bind the human tracheal section. The Q226L mutant, on the other hand, failed to bind at all, similarly as observed in the glycan array, yet a tissue section would display many more different types of glycans compared to our quite restricted glycan array. The H5 protein from Akita16, on the other hand, displays a concentration-dependent binding to both the chicken as well as the human tracheal section, with some binding to the human trachea at high concentration. This tissue image was reversed entirely when the Q226L mutation was introduced, with only minor binding at high concentration to the chicken trachea and a significant signal at low concentration for the human trachea. France16 derived HA strongly binds to the chicken trachea, in which a plateau has already been reached at 2ug/mL. This wild-type protein surprisingly also bind human tracheal tissue sections at low concentration already, despite not binding human-type receptors. Several reports indicate that 2.3.4.4b HA proteins have a broader receptor binding profile compared to classical H5 proteins (29, 30, 39). This indicates that France16 derived HA apparently binds human tracheal sections due to this reported promiscuity. Indeed, a related 2.3.4.4b H5N1 virus, A/Caspian gull/Netherlands/1/2022, efficiently bound to human upper respiratory tract tissues (40). The introduction of Q226L in the cleavage site modified France16 HA did not induce a significant difference in receptor binding specificity, now with diminished binding to chicken trachea at low concentration but similar binding to human trachea compared to wild type. However at high concentration, the France16 Q226L mutant binds with higher intensity compared to wild-type as one get closer to saturation. Thus, especially in the 2.3.4.4e HA, a single Q226L mutation converts binding from chicken epithelial cells to those of humans in the upper respiratory tract.

### Direct concentration-dependent binding reveals the specificity change in A/black swan/Akita/1/16 due to Q226L

To better understand how the Q226L mutants affect receptor binding strength, we subjected the WT and mutant proteins to a direct receptor binding assay in which we conjugated biotinylated structures on streptavidin-coated plates. Linear triLacNAc structures were coated with sialosides either terminating with α2,3 or α2,6 linked Sia (3SLN3 and 6SLN3, where LN stands for LacNAc, which is galactose β1-4-linked to N-acetylglucosamine), as commonly employed by many laboratories (41, 42). For Indo05 HA, we again observed a loss in binding due to the Q226L mutation, similar to in the glycan array. Also similar to the two previous assays, we observed binding to 6SLN3 with the Q226L mutation in Akita16. The modified H5 protein derived from France16 solely bound avian-type receptors, similar to the Q226L mutant. As in previous assays, we did not observe any human-type receptor binding, except at high concentrations.

### Structural characterization of A/black swan/Akita/1/16 HA and mutant with avian- and human-type receptor analog

To decipher the interaction between HA and ligand, we determined crystal structures of wild type (WT) and Q226L (Glu226Leu) mutant of Akita16 HA with avian- and human-type receptor analogs. The Akita16-WT HA with avian-type receptor analog, LSTa (NeuAcα2-3Galβ1-3GlcNAcβ1-3Galβ1-4Glc), was determined at 2.71 Å resolution (Fig. 4A, Table S1). In this complex, only two monosaccharide moieties of LSTa are well-ordered based on the interpretable electron density (Fig. S1A). The LSTa adopt a canonical *trans* conformation in the glycosidic bond between Sia-1 and Gal-2. The avian signature residue Q226 forms several hydrogen bonds via its side chain with Sia-1 and Gal-2. Other interactions to Sia1, and Gal-2 are from the HA130-loop and 190-helix (Fig. 4A). We then compared the structure of Akita16-WT-LSTa complex with the Indo05-WT-LSTa complex (Fig. 4B). The overall interactions among HA, Sia-1, and Gal-2 in Indo05 are simliar to those in Akita16-WT (Fig. 4C), despite A137 and the 133a deletion in Akita16-WT. However, in Akita16-WT, Gal-2 is slightly closer to 190-helix due to the hydrogen bond between the O-6 of Gal-2 and the side-chain carboxyl of E190 (Fig. 4A).

**Fig. 4.**
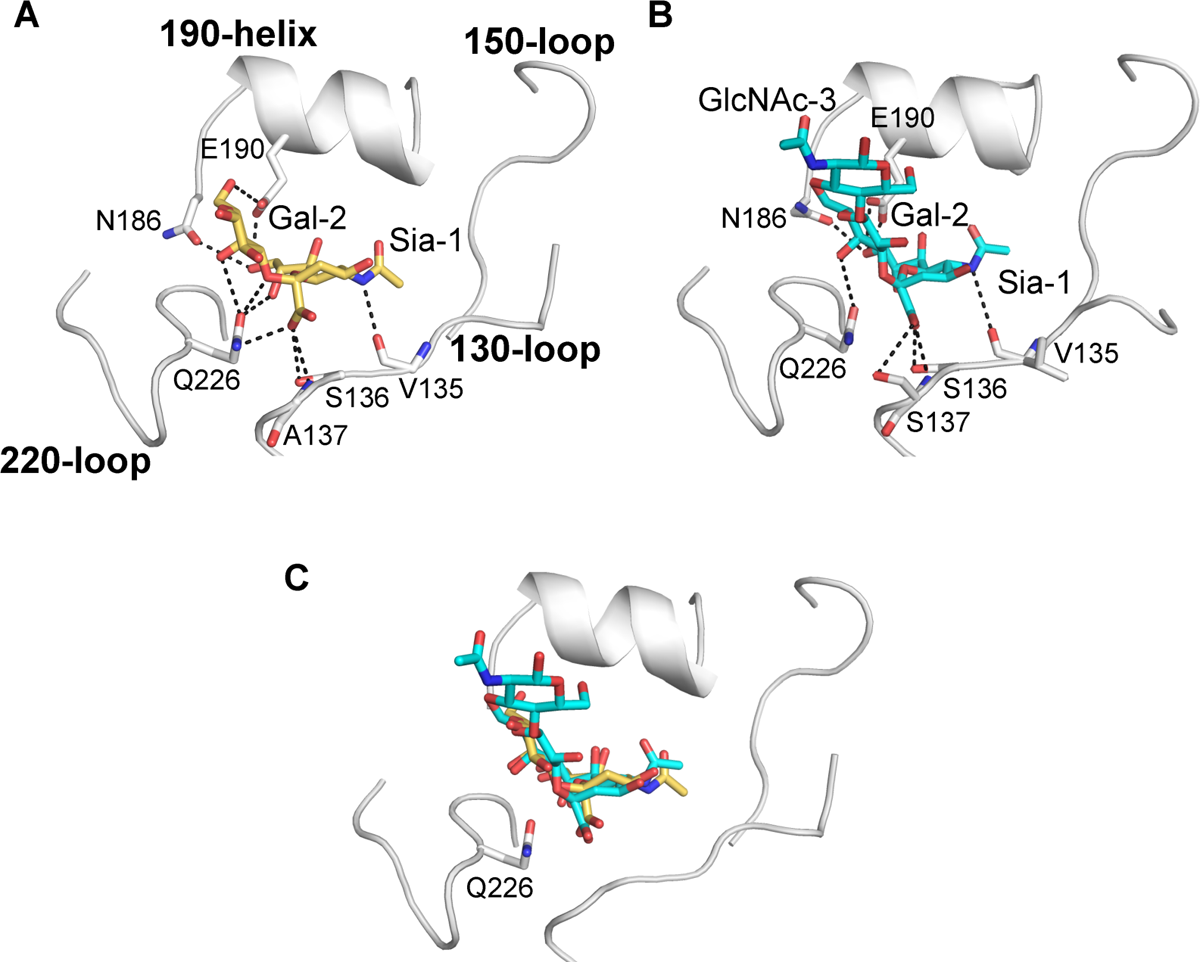
Crystal structure of A/black swan/Akita/1/16 and A/Indonesia/5/2005 H5 HA with avian receptor analog LSTa. The receptor binding site (RBS) is shown as a backbone cartoon. HA residues involved in the interaction with the receptor analog are presented as sticks. Hydrogen bonds are indicated as black dashes. Avian-type receptor analog LSTa (NeuAcα2-3Galβ1-3GlcNAcβ1-3Galβ1-4Glc) is used for both HA. (**A**) Akita16-WT HA with LSTa (**C**). (**B**) Indo05-WT HA with LSTa (PDB: 4K63). (**C**) Superposition of LSTa in Akita16-WT HA (yellow) and Indo05-WT HA (cyan).

The Q226L substitution confered significant receptor binding ability to human-type receptors in Akita16-Q226L mutant, as shown in all receptor bindings assays (Figs.1-3). We therefore determined the crystal structure of Akita16-Q226L mutant with human-type receptor analog LSTc (NeuAcα2-6Galβ1-4GlcNAcβ1-3Galβ1-4Glc) at a resolution of 2.52 Å to decipher the receptor-ligand interaction (Fig. 5A, Table S1). In this structure, three of five monosaccharide moieties are well-ordered (Fig. S1B). The Sia-1 and Gal-2 adopt a *cis* conformation, and the GlcNAc-3 exit the RBS toward 190-helix (Fig. 5A). Residue L226 makes van der Waals’ interactions with the non-polar portions of LSTc (Fig. 5A). As human-type receptor specificity is vital for airborne transmission by respiratory droplets, we compared the LSTc in Akita16-Q226L with a crystal structure of an H5 HA protein (Indo05mut (21)) derived from a mutant Indo05 virus (with four HA mutations Q226L/G228S and (polymerase basic 2) PB2 mutation E627K) that enabled airborne transmission among ferrets (16). The Indo05mut H5 HA demonstrated increased binding to human-type receptors but substantially decreased binding to avian-type receptors (21). In this Indo05 ferret-transmissible Indo05mut HA, the O-4 of Gal-2 hydrogen bonds with the carbonyl oxygen of G225 (Fig. 5B), resulting in LSTc coming closer to 220-loop (Fig. 5C). The orientation of GlcNAc-3 in Indo05mut is rotated ∼30° and seems to move away from the 190-helix due to the longer side chain at R193 (N193 in Akita16 HA) (Fig. 5). This may result in weaker binding to human-type receptors as a bulky side chain at position 193 is detrimental for human-type receptor binding (14).

**Fig. 5.**
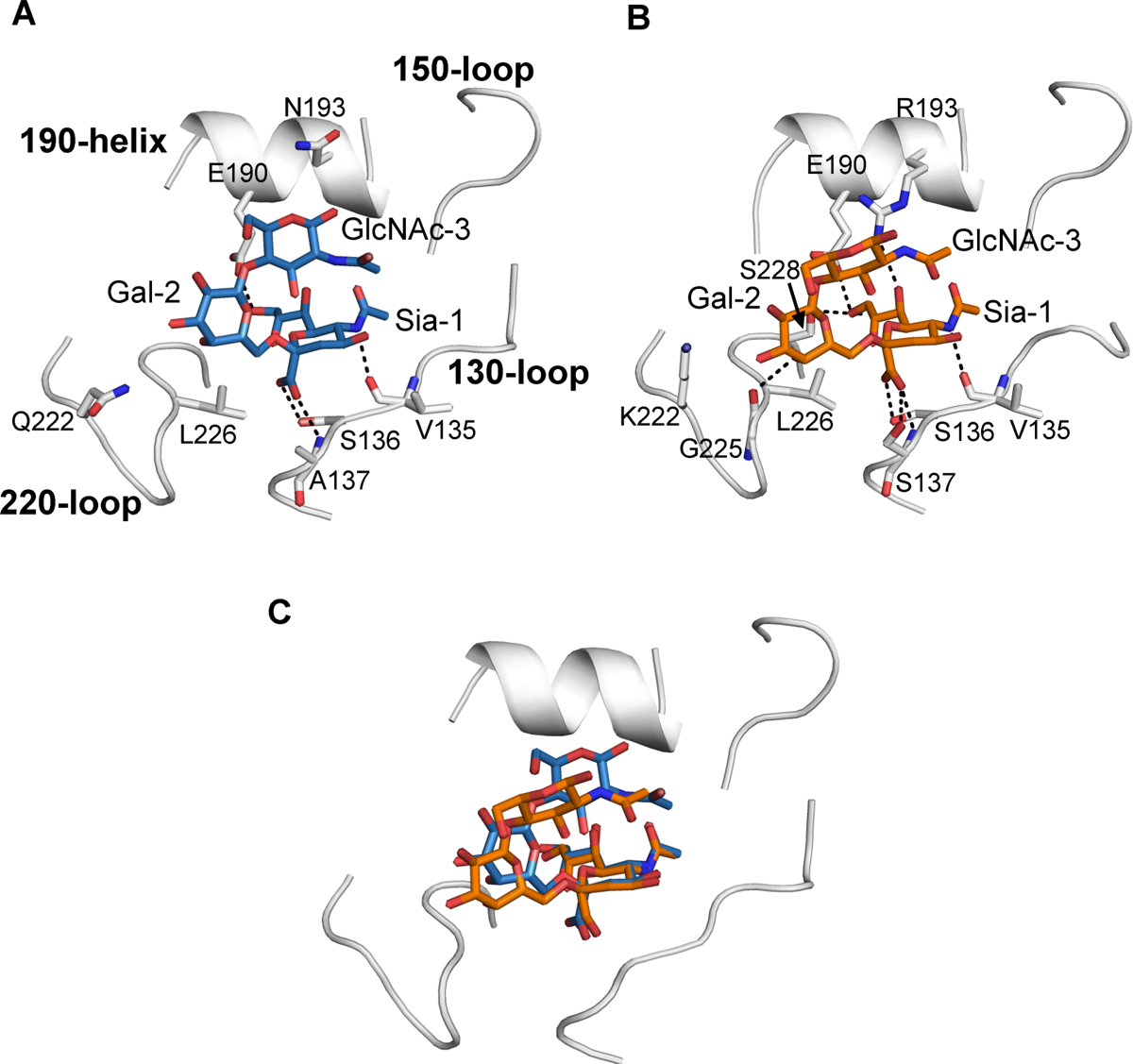
Crystal structure of A/black swan/Akita/1/1utant with human receptor analog LSTc and comparison with other H5 HAs with LSTc. Receptor binding site (RBS) is shown as back bone cartoon. Residues that involved in the interaction between HA and receptor analog are presented as sticks. Hydrogen bonds are indicated as black dashes. LSTc (NeuAcα2-6Galβ1-4GlcNAcβ1-3Galβ1-4Glc) is used as Human-type receptor analog. (**A**) Akita16-Q226L mutant with LSTc. (**B**) Indo05mut HA with LSTc (PDB: 4K67). (**C**) Superposition of LSTc in Akita16-Q226L mutant with Indo05mut HA (orange).

### Structural characterization of A/duck/France/161108h/2016 Q226L mutant with human-type receptor analog

The Q226L substitution in clade 2.3.4.4b France16 HA in our assays increases human-type receptor binding ability but only very slightly, and still retains avian-type receptor binding (Fig. 1-3). To assess the interaction between avian-type receptor and RBS in the France16-Q226L mutant, we determined the crystal structure of France16-Q226L with LSTa at 2.65 Å (Fig. 6A). In this structure, four of the five monosaccharide moieties of LSTa are well-ordered (Fig. S1C). The LSTa still adopts a *trans* conformation in the glycosidic bond between Sia-1 and Gal-2, which is a canonical conformation observed when LSTa binds to avian influenza hemagglutinin (Fig. 6A). Sia-1 forms several hydrogen bonds with residues from the 130-loop and 190-helix, similar to LSTa in Akita16-WT (Fig. 5A and 6A). The orientation of Sia-1 and Gal-2 is similar to the LSTa in Akita16-WT (Fig. 6B). We then compared the France16-Q226L-LSTa complex with human-infecting A/Shanghai/02/2013(H7N9) HA (Sh2), which already contains L226 in its RBS and also retains an avian-type specificity (Fig. 6C) (43). In the Sh2-LSTa complex, only Sia-1 and Gal-2 could be modeled and adopt a *trans* conformation (Fig. 6C) consistent with an avian binding preference (Fig. 6A and 6C) (43).

**Fig. 6.**
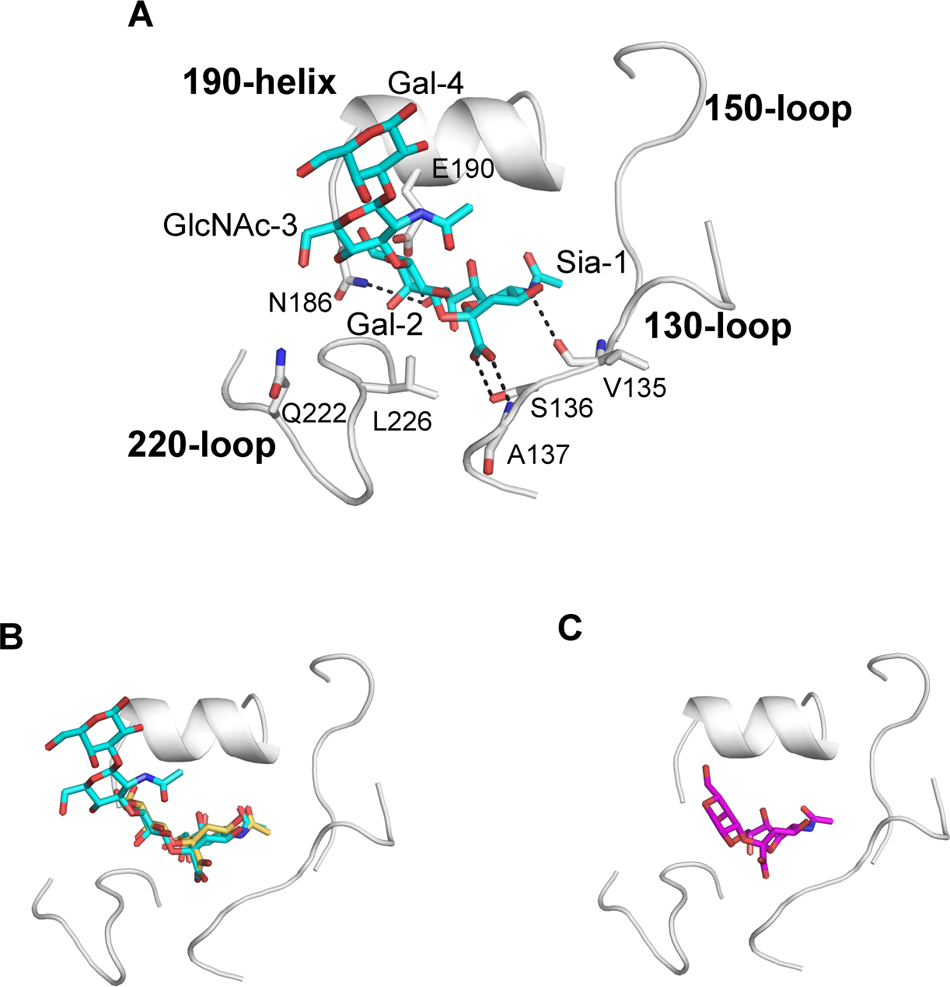
Crystal structure of A/duck/France/161108h/2016 Q226L mutant with avian receptor analog LSTa and the comparison with other H7 HA and H5 HA with LSTa. (A) France16-Q226L mutant with LSTa. (B) Superposition of LSTa in France16-Q226L (cyan) mutant with LSTa in the Akita16-WT (magenta). (C) A/Shanghai/02/2013 (H7N9) HA with LSTa (PDB: 4N5K).

## Discussion

Here, we demonstrate that the Q226L mutation results in different phenotypes in different H5 backbones. Whereas for pre 2.3.4.4 H5 viruses, several additional mutations are necessary (34), in the 2.3.4.4e H5 HA, a single Q226L converts to human-type receptor binding. While 2.3.4.4b H5N1 viruses are currently the dominant strain, the last isolation of an H5N6 2.3.4.4e virus was in 2017 (44, 45).

Our results establish that different genetic and antigenic backgrounds must be studied to determine mutational pathways to human-type receptor binding. Several hallmark mutations have been described that confer avian- to human-type receptor specificity with E190D and G225D for H1 HA and Q226L and G228S for H2 and H3 HA (46). However, the introduction of the H3N2 pandemic came with additional amino acid mutations that affected receptor binding avidity and viral replication (47). Furthermore, these hallmark mutations can not be directly translated by themselves to other subtypes, such as H5 and H7, and multiple amino acid changes are often necessary to change receptor specificity (8, 14, 22, 23). So while single mutations in H5 HA clearly affect receptor binding and cell entry (32), this is not always the case.

It has recently been demonstrated that a single Q226L mutation in a contemporary 2.3.4.4b H5N1 human isolate, A/Texas/324/24, does result in a complete reversion from avian- to human-type receptor specificity (12). This human isolate differs at six amino acids positions (M111L, Q122L, A144T, I199T, A214V, and D240N) from the France16 H5 HA protein used in this study. Furthermore, our France16 H5 protein has the multibasic cleavage site modified to a monobasic cleavage site; a reverse genetic variant was then employed and expressed as HA0. However, no published studies to date have demonstrated that changes in the multibasic cleavage site affect receptor binding, but that needs further study.

Other important differences between our study here and Lin et al. 2024 (12) is the use of SPR-based analyses and the use of complex binantennary N-glycans in ELISA assays. The Q226L containing H5 derived from A/Texas/324/24 HA binds more strongly to longer biantennary glycans. These structures are present in our array and we do observe some low response in Figure 1c to the longer binantennary glycans. Symmetrical biantennary N-glycans have been previously reported to be important for avian-like HAs engineered to bind human-type receptors (14, 22). These structures have been shown to be present in the human upper respiratory tract (38). We recently probed human respiratory tissues with human H3 HA proteins from 2013-2021 and correlated human tracheal binding with engaging asymetrical N-glycans displaying one arm with a human-type sialic acid (48) that further demonstrates the context dependence of analyzing mutations in the HA either acquired from natural evolution or in the laboratory. Nevertheless, in this particular study, we emphasize the strong binding of Akita16 Q226L hemagglutinin to a linear structure displaying a human-type sialic acid (Fig 3). This phenotypic variation among hemagglutinins emphasizes once again that different genetic or antigenic strains must be evaluated at the molecular level to determine which amino acid changes confer human-type receptor binding.

Human-type receptor specificity is only one of the phenotypic changes necessary to transmit between humans. NA activity, HA stability, fusion pH, and polymerase adaptations are also important (11–13), and indeed, H5 viruses have previously been adapted to be transmitted between ferrets (16, 17). As with receptor specificity mutants, mutations confer stability depending on the protein background (49–51). Mammalian adaptations in PB2 E627K and D701N (52) can confer replication in mammalian cells. These are intermittently found to be present during surveillance (53) or are quickly introduced when transmitted to mammals (54–56). Although transmission by direct contact or fomites has been shown to be efficient between ferrets, transmission by respiratory droplets, which emulates human sneezing; was not (39, 56–61). Some human-type receptor binding is found in 2.3.4.4 H5Nx viruses (51, 62, 63); however, avian-type receptor binding was also maintained, and complete conversion is needed for human-to-human transmission (13, 20, 21). Nevertheless, recently an H9N2 bat influenza A virus was shown to be able to transmit between ferrets by respiratory droplets, while lacking observable human-type receptor binding (64). It is important to note that only recombinant HA proteins were used in this study and no viruses were tested or generated.

The France16 HA already displayed binding to human tracheal tissues, despite not binding human-type receptors. Recently, it was demonstrated that a related 2.3.4.4b H5N1 virus, A/Caspian gull/Netherlands/1/2022, bound efficiently to human upper respiratory tract tissues (40). It has been reported that 2.3.4.4b virus HA proteins that circulate in the US bind more promiscuously to avian-type receptors that include a fucose linked to the N-acetylglucosamine, creating sialyl-lewisX (sLeX) structures (30). sLeX has been shown to be specific receptors for some H5 viruses (65, 66); however the presence of sLeX on human tracheal epithelial cells differs between studies and probably depends on the lectin used (67, 68). Thus, the involvement of sLeX in binding human tracheal tissues for influenza A virus HA remains to be elucidated.

With the continuous and massive circulation of 2.3.4.4b viruses in both avian and mammalian species, it may be just a matter of time before different adapted gene segments collide. It is, therefore, vital to know which mutations in NA and HA are essential to adapt to the respiratory tract of humans. One such phenotype, human-type receptor binding, is described here, and we demonstrate that previously circulating 2.3.4.4e viruses only needed a single Q226L mutation to do so. The France16 2.3.4.4b H5N8 HA used in this study has six mutational differencers and a modified cleavage site compared to the A/Texas/37/2024 2.3.4.4b HA (12). Perhaps France16 HA’s ability to bind human tracheal epithelial cells is already an indicator for its zoonotic abilities (40). Also, whether or not the current H5N1 2.3.4.4b HAs would require additional amino acid changes to bind more strongly to less complex glycan structures that are more abundant in the human respiratory tract (38, 69), although increased avidity to complex biantennary glycans may suffice.

## Materials and methods

### Expression and purification of trimeric influenza A hemagglutinins for binding studies

Recombinant trimeric IAV hemagglutinin ectodomain proteins (HA); A/Indonesia/5/05 (34), A/duck/France/1611008h/16 (EPI869809) with the multibasic cleavage site replaced with a monobasic cleavage site (70), and A/black swan/Akita/1/2016 (EPI888450) were cloned into the pCD5 expression vector (addgene plasmid #182546 (71)) in frame with a GCN4 trimerization motif (KQIEDKIEEIESKQKKIENEIARIKK), a mOrange2 (72) and the Twin-Strep-tag (WSHPQFEKGGGSGGGSWSHPQFEK); IBA, Germany). Mutations in HAs were generated by site-directed mutagenesis. The trimeric HAs were expressed in HEK293S GnTI(-) cells with polyethyleneimine I (PEI) in a 1:8 ratio (µg DNA:µg PEI) for the HAs as previously described (73). The transfection mix was replaced after 6 hours by 293 SFM II suspension medium (Invitrogen, 11686029), supplemented with sodium bicarbonate (3.7 g/L), Primatone RL-UF (3.0 g/L, Kerry, NY, USA), glucose (2.0 g/L), glutaMAX (1%, Gibco), valproic acid (0.4 g/L) and DMSO (1.5%). According to the manufacturer’s instructions, culture supernatants were harvested 5 days post-transfection and purified as HA0 with sepharose strep-tactin beads (IBA Life Sciences, Germany).

### Glycan microarray binding studies

HAs (50 µg/ml) were pre-complexed with human anti-strep tag and goat anti-human-Alexa555 (#A21433, Thermo Fisher Scientific) antibodies in a 4:2:1 molar ratio respectively in 50 µL PBS with 0.1% Tween-20. The mixtures were incubated on ice for 15 minutes and then on the array’s surface for 90 minutes in a humidified chamber. Then, slides were rinsed successively with PBS-T (0.1% Tween-20), PBS, and deionized water. The arrays were dried by centrifugation and immediately scanned as described previously (74). The six replicates were processed by removing the highest and lowest replicates and calculating the mean value and standard deviation over the four remaining replicates.

### Protein histochemical tissue staining

Sections of formalin-fixed, paraffin-embedded chicken (*Gallus gallus domesticus*) were obtained from the Division of Pathology, Department of Biomolecular Health Sciences, Faculty of Veterinary Medicine of Utrecht University, the Netherlands. Sections of formalin-fixed, paraffin-embedded human tissues were obtained from the UMC Utrecht, Department of Pathology, Utrecht, the Netherlands (TCBio-number 22-599). In the figures, representative images of at least two individual experiments are shown. Protein histochemistry was performed as previously described (75, 76). In short, tissue sections of 4 µm were deparaffinized and rehydrated, after which antigens were retrieved by heating the slides in 10 mM sodium citrate (pH 6.0) for 10 min. Endogenous peroxidase was inactivated using 1% hydrogen peroxide in MeOH for 30 min at RT. When a neuraminidase treatment was performed, slides were incubated overnight at 37°C with neuraminidase from *Vibrio cholerae* (#11080725001, Roche) diluted 1:50 in a solution of 10mM potassium acetate and 0.1% Triton X-100 at pH 4.2. Non-treated slides in experiments with neuraminidase were incubated with only buffer. Tissues were blocked at 4°C using 3% BSA (w/v) in PBS for at least 90 minutes. For plant lectin stains, blocking was performed using carbo-free blocking solution (SP-5040-125; Vector Laboratories, Burlingame, CA, USA) instead. Subsequently, slides were stained for 90 minutes using anti-sLe^x^ antibody, or pre-complexed HAs as previously described for the glycan microarray. For A/duck/France/161108h/2016, 2 µg/ml of H5 HA were used, while for the other H5 HAs, 2.5 µg/ml was used. For A/NL/109/2003, 5.0 µg/ml H3 HA were used. For antibody control images, only antibodies and no HAs been used.

### Dose-dependent direct binding

Maxisorp plates were coated with 50μl of 5μg/mL streptavidin in PBS overnight at 4°C. After a protein block of 1%BSA in 0.1%PBStween20 for 3 hrs at room temperature, each well was incubated with 50 μl of 50nM solution of biotinylated glycans (3SLN^3^, 6SLN**^3^**) in PBS overnight at 4°C. The plate was subsequently washed with PBS to remove excess glycan and blocked with 1% BSA in 0.1% PBStween20 for 3 hours at room temperature. 20μg/ml strep-tagged HA protein, primary (human anti streptag IgG), and secondary (HRP-conjugated goat anti-human IgG) antibodies were mixed in the ratio 4:2:1 and incubated on ice for 20 min. The mixture (precomplexed HA) was made up to a final volume of 300 μl with PBStween20 buffer, and 50 μl of precomplexed HA was added to each glycan-coated well and incubated at room temperature for 1hr. The wells were extensively washed with PBS containing 0.05% Tween-20 followed by washes with PBS.

### Expression and purification of trimeric influenza A hemagglutinins for structural studies

The HAs for crystal structures were expressed in Hi five cells as described previously (18). Recombinant baculovirus was incorporated with HA gene which contains a GP67 signal peptide at the N-terminus and trimerization domain at C-terminus followed by a thrombin cleavage site and 6x-His tag. The recombinant baculoviruses were generated using a bac-to-bac insect cell expression (Thermo Fisher Scientific). The HAs were harvested 72 hours post-infection from supernatant by metal-affinity chromatography using Ni beads. The purified HA proteins were buffer-exchanged into Tris-buffered saline (TBS) (pH 8), digested with trypsin in a final ratio of 1:1000 (wt/wt) and then further purified by size exclusion chromatography (SEC) using Hiload 16/90 Superdex 200 column (GE healthcare).

### Crystallization, data collection, and structure determination

The purified soluble WT HA and mutant were concentrated to 10mg/ml for crystallization screening. The initial crystallization conditions for WT HA and mutant were achieved from crystal screening via our automated Rigaku CrystalMation system at The Scripps Research Institute using JCSG Core Suite (QIAGEN) as precipitant. The crystallization screening was set up according to the sitting drop vapor diffusion method by mixing 0.1 μl of protein with 0.1 μl of the reservoir solution. The optimized crystallization conditions were: 0.2M sodium iodide, 20% PEG 3350 for Akita16-WT, 0.2M ammonium format, 20% PEG3350 for Akita16-Q226L, and 0.1M Tris, 2M ammonium sulfate, 0.2M lithium sulfate for France16-Q226L. For HA-receptor analog complexes, avian-type receptor analog LSTa (Neu5Acα2-3Galβ1-3GlcNAcβ1-3Galβ1-4Glc), and human-type receptor analog LSTc (Neu5Acα2-6Galβ1-3GlcNAcβ1-3Galβ1-4Glc) were used. Apo-crystals for each HA were harvested after 10 days and soaked with ligands at a final concentration of 5mM in the reservoir solution with 10-15% 10% (v/v) ethylene glycol for 5 minutes. The crystals were then stored in liquid nitrogen until data collection. Diffraction data for Apo-HA and HA-ligand complexes were collected at synchrotron radiation beamlines and integrated and scaled using HKL2000 (77) (Table S1). Initial phases for Apo-HA and HA-ligand complexes were solved by molecular replacement using Phaser with H5 HA (PDB: 5E30) as a model (78). Model bulding and refinement were carried out by Coot and Phenix (79, 80) (Table S1).

## Acknowledgements

R.P.d.V. is a recipient of an ERC Starting grant from the European Commission (802780) and is supported by the Mizutani Foundation for Glycoscience. This research was made possible by funding from ICRAD, an ERA-NET co-funded under the European Union’s Horizon 2020 research and innovation programme (https://ec.europa.eu/programmes/horizon2020/en), under Grant Agreement n°862605 (Flu-Switch) to R.P.d.V. R.P.d.V and M.R.C are supported by an NWO-M2 (OCENW.M20.106). The glycan array setup was supported by the Netherlands Organization for Scientific Research (NWO, TOP-PUNT 718.015.003 to G.-J. B.). This structural biology research at Scripps Research was partially funded by National Institutes of Health NIAID Centers of Excellence for Influenza Research and Response contract 75N93021C00015 / PENN CEIRR (I.A.W.). X-ray diffraction datasets were collected at the Stanford Synchrotron Radiation Lightsource (SSRL) beamline 12-1. Use of the Stanford Synchrotron Radiation Lightsource, SLAC National Accelerator Laboratory, is supported by the U.S. Department of Energy, Office of Science, Office of Basic Energy Sciences under Contract No. DE-AC02-76SF00515. The SSRL Structural Molecular Biology Program is supported by the DOE Office of Biological and Environmental Research, and by the National Institutes of Health, National Institute of General Medical Sciences (P30GM133894). The contents of this publication are solely the responsibility of the authors and do not necessarily represent the official views of NIGMS or NIH. We gratefully acknowledge all data contributors, i.e., the authors and their originating laboratories responsible for obtaining the specimens and their submitting laboratories for generating the genetic sequence and metadata and sharing via the GISAID Initiative, on which part of this research is based.

---

**Table S1.** X-ray data collection and refinement statistics.

| **Data collection** | Akita16 | Akita16-Q226L | France16-Q226L |
| --- | --- | --- | --- |
|  | +LSTa | +LSTc | +LSTa |
| Beamline | SSRL12-1 | SSRL12-1 | SSRL12-1 |
| Wavelength (Å) | 0.9795 | 0.9795 | 0.9795 |
| Space group | C2 | C2 | P2<sub>1</sub>2<sub>1</sub>2<sub>1</sub> |
| Unit cell parameters |  |  |  |
| a, b, c (Å) | 187.2 100.4 132.1 | 188.1 99.8 132.1 | 95.9 176.4 226.4 |
| α, β, γ (°) | 90 125.9 90 | 96, 126.2, 90 | 90, 90, 90 |
| Resolution (Å)<sup>a</sup> | 50.0-2.71 (2.76-2.71) | 50.0-2.52 (2.56-2.52) | 50.0-2.65 (2.70-2.65) |
| Unique reflections<sup>a</sup> | 52,384 (5206) | 66,157 (6297) | 110,745 (10,532) |
| Redundancy<sup>a</sup> | 6.2 (5.9) | 6.8 (6.8) | 11.4 (11.3) |
| Completeness (%)<sup>a</sup> | 98 (100) | 98 (94) | 100 (100) |
| <I/σ><sup>a</sup> | 13.2 (1.2) | 23.8 (1.8) | 10.8 (0.6) |
| R<sub>sym</sub><sup>b</sup> (%)<sup>a</sup> | 21.6 (>100) | 11.7 (88) | 27.4 (>100) |
| R<sub>pim</sub><sup>b</sup> (%)<sup>a</sup> | 9.6 (63) | 4.9 (36) | 8.5 (85) |
| CC<sub>1/2</sub><sup>c</sup> (%)<sup>a</sup> | 98 (63) | 99 (78) | 99 (39) |
| **Refinement statistics** |  |  |  |
| Resolution (Å) | 35.7-2.71 | 39.68-2.52 | 26.1-2.65 |
| Reflections (work) | 52,313 | 65,129 | 110,720 |
| Reflections (test) | 5197 | 6297 | 10,528 |
| R<sub>cryst</sub><sup>d</sup> / R<sub>free</sub><sup>e</sup> (%) | 21.8/26.9 | 19.9/25.6 | 20.3/23.5 |
| No. of copies in ASU | 3 | 3 | 3 |
| No. of atoms | 12,069 | 12,294 | 12,173 |
| HA | 119,11 | 11,893 | 11,825 |
| glycan | 109 | 207 | 229 |
| Receptor analog | 32 | 138 | 57 |
| Solvent | 17 | 56 | 62 |
| Average B-values (Å<sup>2</sup>) | 70 | 60 | 52.8 |
| HA | 69 | 59 | 73 |
| glycan | 95 | 92 | 113 |
| Receptor analog | 98 | 100 | 103 |
| Solvent | 53 | 51 | 66 |
| Wilson B-value (Å<sup>2</sup>) | 64 | 59 | 67 |
| **RMSD from ideal geometry** |  |  |  |
| Bond length (Å) | 0.004 | 0.003 | 0.003 |
| Bond angle (°) | 0.58 | 0.57 | 0.61 |
| **Ramachandran statistics (%)<sup>f</sup>** |  |  |  |
| Favored | 96.5 | 97.1 | 97.3 |
| Outliers | 0.0 | 0.2 | 0.0 |
| **PDB code** | pending | pending | pending |

<sup>a</sup> Numbers in parentheses refer to the highest resolution shell.
<sup>b</sup> $R_{\text{sym}} = \sum_{hkl} \sum_i |I_{hkl,i} - \langle I_{hkl} \rangle| / \sum_{hkl} \sum_i I_{hkl,i}$ and $R_{\text{pim}} = \sum_{hkl} (1/(n-1))^{1/2} \sum_i |I_{hkl,i} - \langle I_{hkl} \rangle| / \sum_{hkl} \sum_i I_{hkl,i}$ , where $I_{hkl,i}$ is the scaled intensity of the $i^{\text{th}}$ measurement of reflection $h, k, l$ , $\langle I_{hkl} \rangle$ is the average intensity for that reflection, and $n$ is the redundancy.
<sup>c</sup> CC<sub>1/2</sub> = Pearson correlation coefficient between two random half datasets.
<sup>d</sup> $R_{\text{cryst}} = \sum_{hkl} |F_o - F_c| / \sum_{hkl} |F_o| \times 100$ , where $F_o$ and $F_c$ are the observed and calculated structure factors, respectively.
<sup>e</sup> $R_{\text{free}}$ was calculated as for $R_{\text{cryst}}$ , but on a test set comprising 5% or 10% of the data excluded from refinement.
<sup>f</sup> From MolProbity (81).

**Fig. S1.**
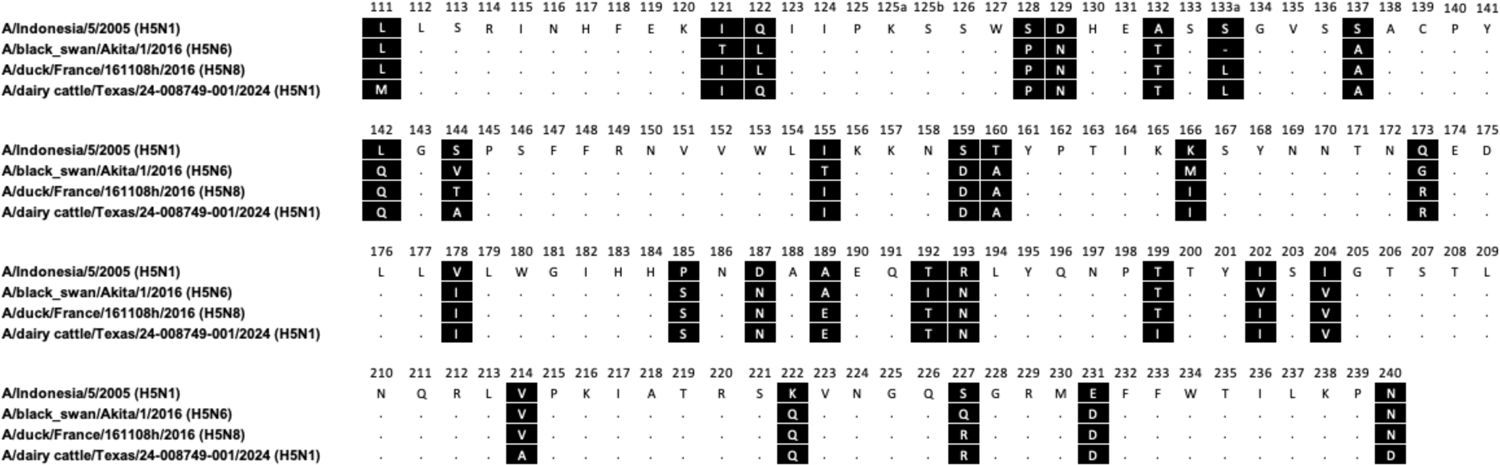
Amino acid aligment of the H5 protein used in the study. Added sequence is the first bovine sequence for comparison to Akita16 and France16 respectively

**Fig. S2.**
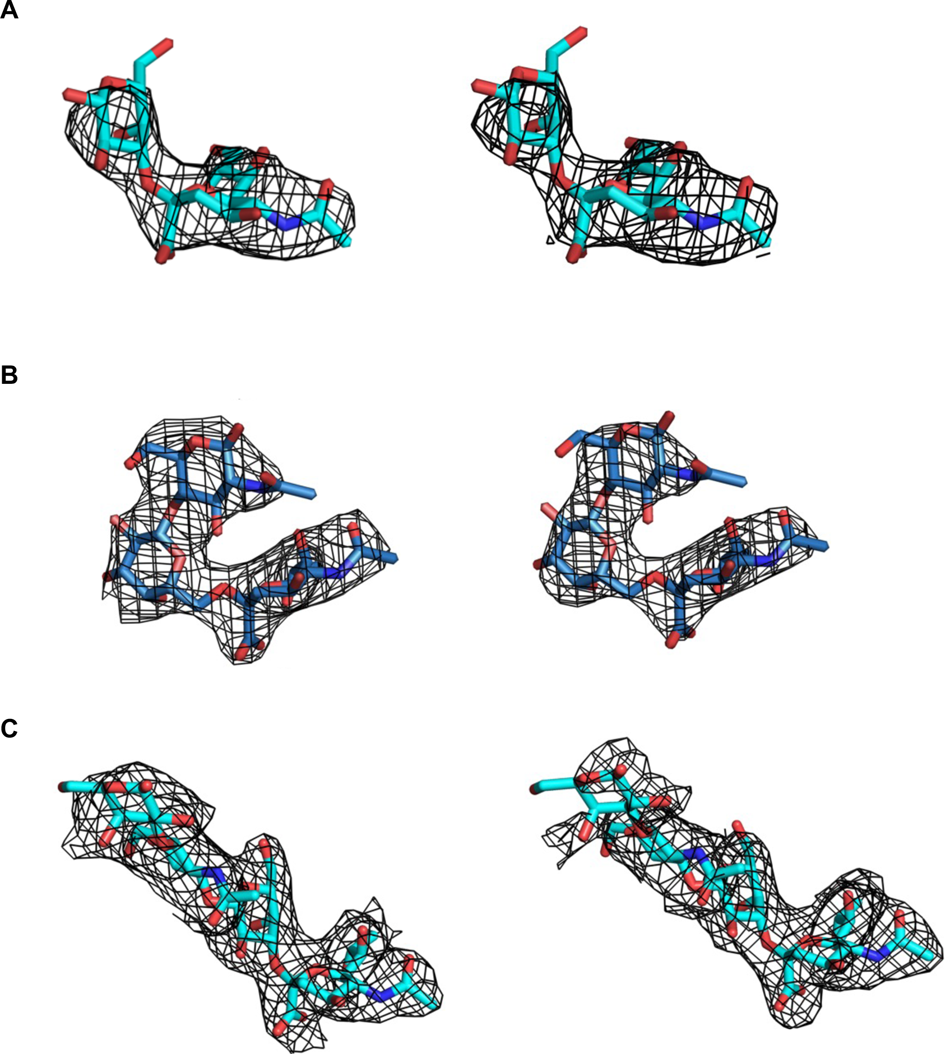
Electron density maps of avian and human receptor analogs in HA-ligand complexes. 2Fo-Fc (left panel) and unbiased omit (right panel). Avian receptor LSTa (NeuAcα2-3Galβ1-3GlcNAcβ1-3Galβ1-4Glc) is shown as cyan. Human receptor analog LSTc (NeuAcα2-6Galβ1-4GlcNAcβ1-3Galβ1-4Glc) is represented as skyblue. Electron density maps for each ligand are contoured at the 1 σ level and represented in a mesh with its refined structure superimposed, respectively. (**A**) LSTa in Akita16-WT HA. (**B**) LSTc in Akita16-Q226L mutant. (**C**) LSTa in France16-Q226L mutant.

